# Sex-specific transcriptional signatures of oxycodone persist during withdrawal and abstinence in the suprachiasmatic nucleus of heterogeneous stock rats

**DOI:** 10.1101/2025.04.29.651331

**Authors:** Tara C. Delorme, Snehal Sambare, Benjamin R. Williams, Mackenzie C. Gamble, Leah C Solberg Woods, Lisa Maturin, Abraham A. Palmer, Olivier George, Ryan W. Logan

**Affiliations:** Department of Psychiatry, University of Massachusetts Chan Medical School, Worcester, MA, USA; Department of Pharmacology, Physiology & Biophysics, Boston University School of Medicine, Boston, MA, USA; Department of Internal Medicine, Section on Molecular Medicine, Wake Forest University School of Medicine, Winston-Salem, NC, USA; Department of Psychiatry, University of California, San Diego, La Jolla, CA, USA; Department of Neurobiology, University of Massachusetts Chan Medical School, Worcester, MA, USA

**Keywords:** opioid use disorder, circadian rhythms, suprachiasmatic nucleus, RNA sequencing, sex differences, heterogeneous stock rats, intravenous self-administration

## Abstract

Opioid use disorder (OUD) is a major public health issue. Sleep and circadian disruptions are recognized as hallmarks of opioid addiction, often emerging during withdrawal and lasting into abstinence. However, little is known about the impact of opioids on the brain’s primary circadian pacemaker, the suprachiasmatic nucleus (SCN). We examined SCN transcriptomic changes in genetically diverse heterogeneous stock rats across different opioid physiological and behavioral states (naïve, oxycodone intoxication, acute withdrawal, and prolonged abstinence), alongside behavioral assessments. In females, intoxication and withdrawal altered pathways related to neurotransmission, circadian rhythms, and inflammation, while in males, changes involved immune regulation, DNA damage, and metabolism. During abstinence, females showed enrichment in stress-related pathways, particularly those involved in energy metabolism and neurotransmitter function, whereas males exhibited enrichment in pathways related to cellular detoxification and oxidative stress, suggesting lasting, sex-specific effects of oxycodone administration during withdrawal and abstinence. Further, the highest proportion of sex-specific rhythmic differentially expressed genes (DEGs) were identified during abstinence compared to other states, suggesting sex differences in gene expression in the SCN during opioid abstinence. Co-expression network analysis identified a black module linked to synaptic signaling and a red module linked to ciliary function, which were positively and negatively associated with intoxication, respectively. Black module genes were positively correlated with addiction-related behaviors during abstinence, while red module genes inversely correlated with these behaviors during intoxication, linking opioid-induced alterations in the SCN to addiction-like phenotypes. These findings highlight the SCN as a dynamic, sex-specific target of opioid exposure and suggests that SCN alterations may contribute to long-term behavioral and physiological consequences of OUD.

**Highlights:**

1. Distinct sex specific SCN gene patterns across opioid physiological and behavior
2. Intoxication in females increased synaptic, glutamatergic, and addiction pathways
3. Circadian entrainment pathway enriched in females after intoxication
4. Rhythmic DE genes may drive sex differences in abstinence
5. SCN gene expression correlated with addiction-like behaviors

## 1. Introduction

Opioid use disorder (OUD) remains a major public health crisis, affecting approximately 7.6 million people in the U.S., representing nearly half of global cases [1]. The opioid epidemic traces back to the 1990s, when increased emphasis on pain management led to a surge in opioid prescriptions, particularly oxycodone [2]. Oxycodone, a potent and highly addictive painkiller, played a central role in this crisis [3, 4]. By 2016, Canada and the U.S. had prescribed over 440 million opioids [5, 6]. According to the National Institute on Drug Abuse, approximately 12 percent of individuals prescribed opioids are at risk of developing OUD [7], and 75% of individuals with OUD report their misuse began with prescription opioids [8]. A SAMHSA survey further estimates that over 2.1 million Americans live with OUD involving prescription opioids alone [9]. The impact of opioid misuse is staggering, with opioids accounting for over half of all drug-related deaths in 2017 [10]. These data highlight the urgent need to study opioids— including oxycodone—as part of ongoing efforts to address OUD.

One promising avenue for improving treatment outcomes in opioid use disorder (OUD) involves targeting disruptions in sleep and circadian rhythms, which are increasingly recognized as both causes and consequences of opioid addiction. An estimated 60–70% of individuals with OUD meet criteria for circadian-related disorders, often experiencing poor sleep quality and disrupted hormonal rhythms, including altered corticosterone and melatonin cycles [11–14]. Chronic opioid use worsens these disturbances [15, 16], while circadian misalignment can increase cravings and negative affect, elevating relapse risk [17, 18]. These disruptions may persist for months into abstinence, contributing to the high relapse rates observed in OUD [19–21]. In adults without OUD, acute opioid administration has been shown to reduce stage 2 non-rapid eye movement sleep and rapid eye movement sleep [22], while individuals receiving OUD treatment often exhibit chronically altered rest-activity rhythms, suggesting impaired circadian regulation [23]. Our laboratory found that postmortem brain tissue from individuals with OUD exhibited disrupted molecular and synaptic proteome rhythms, indicating that opioid exposure perturbs the endogenous molecular clock [24, 25]. In rodent models, drug-seeking behavior varies across the circadian cycle and is influenced by constant light exposure [26, 27], and opioid-induced circadian disruptions can be partially rescued by melatonin treatment [28]. Furthermore, we showed that male and female mice deficient in the core circadian gene *Neuronal PAS Domain Protein 2* show altered opioid tolerance and withdrawal symptoms [25]. Together, these findings demonstrate that opioid exposure disrupts circadian rhythms at the transcriptional, translational, and behavioral levels in both humans and animals. Despite this, the molecular basis of circadian dysfunction in OUD remains poorly understood, particularly within the suprachiasmatic nucleus (SCN)—the brain’s master circadian clock—and it is unclear how SCN transcriptomic alterations during key phases of opioid exposure, including withdrawal and abstinence, relate to addiction-relevant behaviors.

In the present study, we examine transcriptomic changes in the SCN of male and female heterogeneous stock (HS) rats during four key phases: naïve, intoxication, acute withdrawal, and prolonged abstinence from intravenous oxycodone self-administration. Compared to other studies which use inbred rodent strains, we leverage the genetic variability of HS rats to more closely mimic the genomic diversity found across humans with OUD. This work aims to uncover sex-specific molecular mechanisms within the SCN that contribute to OUD, with the goal of informing more targeted and effective treatments.

## 2. Methods

### 2.1. Rats

Heterogeneous stock (HS) rats (Rat Genome Database NMcwiWFsm #13673907), bred to maximize genetic diversity by outcrossing eight inbred rat strains (ACI/N, BN/SsN, BUF/N, F344/N, M520/N, MR/N, WKY/N, and WN/N), were supplied by Dr. Leah Solberg Woods (formerly at the Medical College of Wisconsin, currently at Wake Forest University School of Medicine; n = 539). Upon arrival at 3-4 weeks old, rats were quarantined for 2 weeks before being housed in pairs in a temperature (20-22°C) and humidity (45-55%) regulated vivarium in a reversed 12-hour light/dark cycle. Rats had *ad libitum* access to tap water and food pellets (PJ Noyes Company, Lancaster, NH, USA). All experimental procedures were carried out in full compliance to the National Institutes of Health Guide for the Care and Use of Laboratory Animals and were approved by the Institutional Animal Care and Use Committees of The Scripps Research Institute and UC San Diego.

### 2.2. Drug and Intravenous Catheterization

As previously described [29], oxycodone (National Institute on Drug Abuse, Bethesda, MD) was dissolved in 0.9% sterile saline and administered intravenously at a dose of 150 µg/kg/infusion. This dose was chosen based on earlier studies [30, 31] and because it achieves plasma oxycodone levels consistent with those observed in clinical settings (40 ng/ml) [32]. Figure 1 illustrates the experimental design for oxycodone self-administration in rats and the subsequent collection of brain tissue from the SCN for bulk RNA sequencing. As previously described [29], male and female HS rats were anesthetized with vaporized isoflurane (1-5%). Sterile intravenous catheters were surgically implanted into the right jugular vein following a previously established protocol [30]. Rats were given 1 week to recover and were trained to intravenously self-administer oxycodone over an 18-day paradigm consisting of 4 short access, 2-hour self-administration sessions, and 16 long access, 12-hour sessions. Daily monitoring and catheter maintenance were performed by flushing the catheter with heparinized saline (10 U/ml heparin sodium; American Pharmaceutical Partners, Schaumberg, IL, USA) in 0.9% bacteriostatic sodium chloride (Hospira, Lake Forest, IL, USA), containing 52.4 mg/0.2 ml of Cefazolin.

**Figure 1.**
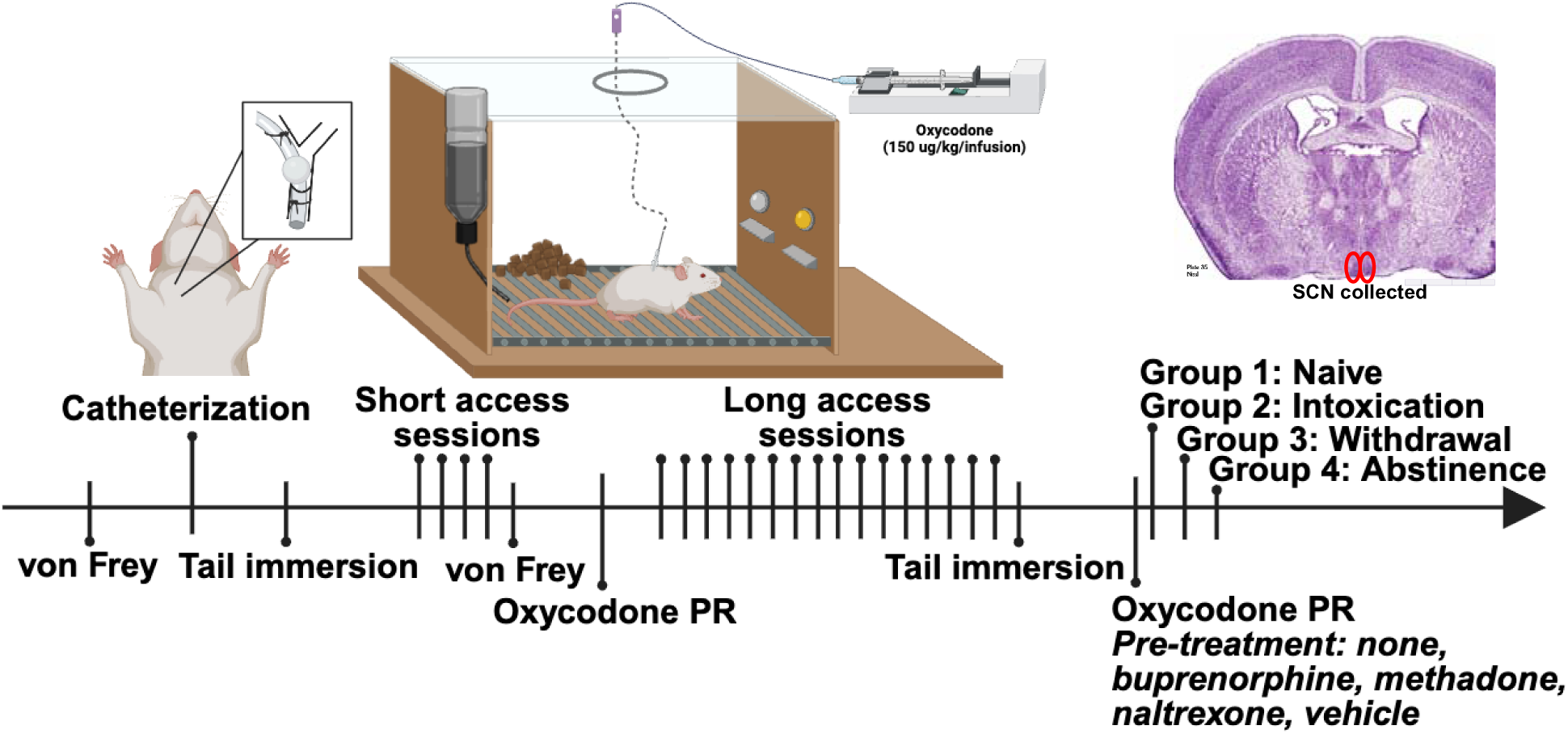
Experimental design for RNA sequencing of the suprachiasmatic nucleus (SCN) in female and male rats following oxycodone self-administration. Rats were implanted with jugular catheters for intravenous self-administration of oxycodone (150 µg/kg/infusion). Following a one-week recovery, rats completed a self-administration protocol consisting of four short-access (2 hour) and sixteen long-access (12 hour) sessions. A battery of behavioral tests was conducted to assess addiction-like phenotypes, including mechanical sensitivity (von Frey), analgesic tolerance (tail immersion), and progressive ratio (motivation) with or without pre-treatment with opioid agonists and antagonists. Rat were sacrificed at group-specific timepoints: naïve (n=6F, 5M; no drug, no surgery), intoxication (n=6F, 5M; 2 h post-session), withdrawal (n=5F, 6M; 12 h post-session), and abstinence (n=6F, 5M; 4–5 weeks post-session). Bilateral SCN punches were collected and processed for bulk RNA sequencing to examine state- and sex-specific transcriptomic changes.

### 2.3. Operant self-administration and other addiction related behaviors

As previously described [29, 33], oxycodone self-administration was conducted in operant conditioning chambers (29 cm × 24 cm × 19.5 cm; Med Associates, St. Albans, VT, USA) housed in sound-attenuating, ventilated cubicles. Each chamber had transparent plastic front doors and back walls, with the remaining walls constructed from opaque metal. Two retractable levers were positioned on the front panel, and both were extended at the start of each session. Oxycodone (150 µg/kg/infusion in saline) was delivered via plastic catheter tubing connected to an infusion pump. Responses on the active lever triggered drug delivery under a fixed-ratio 1 (FR1) schedule, dispensing 0.1mL of oxycodone over 6 seconds. This was followed by a 20-second timeout period, indicated by the illumination of a cue light above the active lever, during which lever presses had no consequence. Responses on the inactive lever were also recorded but had no scheduled effect. Operant responding during timeout periods was logged, and composite metrics for escalation of intake were calculated. A computer running MED-PC IV software-controlled fluid delivery and logged behavioral data. Rats first underwent short access training consisting of four 2-hour sessions over one week. This was followed by long access sessions over six weeks, consisting of 16 total sessions (12 hours/day, 5 days/week). Sessions began at the onset of the dark phase of the light/dark cycle, and food was available during sessions. To assess addiction-like phenotypes, a battery of behavioral tests was administered outlined below.

#### 2.3.1. Progressive ratio

Motivation for oxycodone seeking was assessed using a progressive ratio (PR) schedule of reinforcement. PR sessions were conducted following both the short and long access phases of oxycodone self-administration. At the conclusion of behavioral testing, pharmacological challenges were conducted to assess the effects of opioid agonists and antagonists on motivation for oxycodone. These challenges, administered prior to PR sessions, included buprenorphine (0.5 mg/kg), methadone (3 mg/kg), naltrexone (3 mg/kg), and vehicle, delivered in a within-subject Latin square design. Each drug challenge session was separated by a non-treatment long access session to avoid carryover effects. The response requirement increased according to a modified exponential progression: 1, 1, 2, 2, 3, 3, 4, 4, 5, 5, 6, 6, 7, 7, 8, 8, 9, 9, 10, 10, 11, 12, 13, 14, and so on. The breakpoint was defined as the highest ratio completed before a 60-minute period without a completed response requirement, at which point the session was terminated automatically.

#### 2.3.2. Von Frey test

Mechanical sensitivity was assessed using an electronic von Frey device (Dynamic Plantar Aesthesiometer, Ugo Basile). The device’s needle was applied to the plantar surface of each hind paw, gradually increasing force from 0 to 40 grams over 20 seconds. The force and latency required to elicit paw withdrawal were recorded. Each hind paw was tested three times, and values were averaged. Testing occurred at baseline (prior to surgery) and 12 hours after the final short access session. Force refers to the average withdrawal force (grams), while time refers to the average latency to paw withdrawal (seconds).

#### 2.3.3. Tail immersion test

To assess opioid-induced analgesia, a tail immersion test was conducted following oxycodone administration. An oxycodone injection (0.15 mg/kg) was delivered intravenously (i.v.). Rats were then restrained in a towel, and the distal portion of the tail was immersed in a 52°C water bath. The latency to withdraw the tail was recorded to assess analgesic tolerance. Tests were conducted before the first short access session and before the last long access session.

### 2.4. Tissue collection and pre-processing of data

In collaboration with the National Institute on Drug Abuse Center of Excellence for Genetics, Genomics, and Epigenetics of Substance Use Disorders in Outbred Rats, we obtained brain samples for RNA-seq from male and female HS rats collected at different stages of oxycodone self-administration (U01DA044451: Dr. Olivier George). Separate cohorts of 44 rats (23 females, 21 males) were euthanized by decapitation while intoxicated (2-hours post final self-administration session; F=6, M=5), during acute withdrawal (12-hours post final self-administration session; F=6, M=6), or during drug abstinence (4-5 weeks post final self-administration session; F=6, M=5). A separate cohort of untreated, drug-naïve rats (F=5, M=5) served as a comparison group. The SCN was micropunched from a frozen brain tissue section (1 mm thick punch, -0.9 mm to -1.9 mm relate to bregma) and processed for next generation bulk RNA sequencing. Pre-alignment quality control was performed at the sequencing platform using FastQC. Alignment was performed using STAR version 2.7.1a to the rat genome assembly rn7, incorporating gene annotations based on mouse orthologs to leverage the more comprehensive mouse transcriptome. Transcripts with at least one count per million (CPM) in 50% of samples were retained, in line with the criteria set by Barko et al. (2022) [34]. After filtering, 13,982 genes (∼89% of the total 15,801 transcripts) met the CPM threshold. RNA-sequencing data were deposited in the Gene Expression Omnibus database (GSE289002).

### 2.5. Differential expression, pathway enrichment analysis and threshold free RRHO analysis

DE analysis was conducted using the voom Limma package in R [35]. The analysis used a full model with an interaction term (group x sex) to examine group and sex-specific differences. A p-value threshold of 0.01 and log2 fold change (FC) > ±0.26 (FC > 1.2 or 20% expression change) were used to define DE genes. Comparisons were made in both females and males between naïve and intoxication (accessing acute drug exposure), naïve vs. withdrawal (assessing lasting drug effects), intoxication and withdrawal, and intoxication and abstinence (assessing changes during the transition from drug taking to cessation). Pathway enrichment analyses were conducted using Metascape (https://metascape.org). Rank-rank hypergeometric overlap (RRHO) was used to assess the overlap of gene expression patterns between female and male rats at each phase of opioid exposure using RRHO2 [36]. RRHO is a threshold-free method that identifies overlap between two ranked lists of DE genes, where the ranking is based on the −log10(p value) multiplied by the effect size direction.

#### 2.5.1. Rhythmic gene analysis

All differentially expressed (DE) genes from each comparison were run through CircaDB (http://circadb.hogeneschlab.org/mouse), a database that provides circadian gene expression profiles across various mammalian tissues, including the rodent SCN. Four SCN-specific mouse datasets were utilized: Mouse OST SCN 2014, Mouse Wild Type SCN, and two versions of the Panda 2002 SCN dataset [37, 38]. Polar plots were generated to visualize the circadian phase distribution of DE genes identified as rhythmic.

#### 2.5.2. Weighted gene co-expression network analysis

Weighted gene co-expression network analysis (WGCNA) was conducted to identify co-expression modules and gene networks associated with the different conditions. To construct the co-expression network, raw RNA-seq counts were normalized using the DESeq2 package, employing the variance-stabilizing transformation to stabilize variance across expression levels. The top 5000 most highly variable genes were retained for analysis and a weighted adjacency matrix was constructed using a soft-thresholding power (16). The dynamic tree-cut method was used to identify gene modules (cut-off threshold of 0.05). Modules were defined based on variance across experimental groups and interrelatedness of genes.

#### 2.5.3. Hub gene and module differential connectivity analyses

Hub gene analysis was conducted to identify key regulatory genes within co-expression networks. Gene modules derived from WGCNA were imported into Cytoscape, and ‘degree’ and ‘betweenness centrality’ were calculated using the CentiScaPe plugin. Each gene was classified into one of four categories based on its centrality metrics: hub-bottleneck (high degree and high betweenness), hub, bottleneck, or neither. Classification thresholds were defined as one standard deviation above the mean for each metric. To assess how module connectivity varied across experimental conditions, module differential connectivity (MDC) was conducted by comparing the intramodular connectivity of each gene between groups. Increased coordination of gene expression within a module in one group relative to another reflected a gain in connectivity, while reduced coordination indicates a loss.

#### 2.5.2. Behavioral correlations

To summarize gene expression patterns within each WGCNA module, principal component analysis (PCA) was performed on the normalized expression values of all genes assigned to a given module. The first principal component (PC1), representing the dominant axis of variation within each module, was extracted and used as a module eigengene proxy. PC1 scores were then correlated with behavioral measures using Pearson correlation. Statistical significance was determined using the cor.test function in R, with Benjamini-Hochberg multiple testing correction.

### 2.6. Statistics

Escalation of drug intake was quantified using an escalation index, defined as the ratio of the Z-score for each animal’s intake averaged across the final three days of the long access phase (Zf) to the Z-score on the first day of escalation (Z0) (E = Zf / Z0). Pain index was calculated during withdrawal by expressing each animal’s mechanical threshold as a percentage of their own baseline threshold, which was then converted to a Z-score. Motivation for oxycodone was assessed using a PR index, defined as the Z-score of the breakpoint following long access session. Tolerance to oxycodone-induced analgesia index was calculating the Z-score of the change in tail withdrawal latency following drug administration before the first short access session versus after the final long access session. The addiction index (AI) was calculated by averaging relevant behavioral indices. All statistical analyses were conducted in R (version 4.2.2).

## 3. Results

### 3.1. Transcriptomic changes in female SCN during opioid exposure reveal acute disruptions and sustained stress responses

In female rats, opioid exposure induced widespread transcriptomic changes in the SCN, particularly during intoxication and withdrawal, with sustained molecular alterations persisting into abstinence. Across all comparisons, a total of 3030 DE transcripts were identified, with more genes upregulated (1725) than downregulated (1305). Most DE transcripts (2799) were identified in comparisons with the naïve group (*naïve vs. intoxication*: 1074 DE transcripts, up: 377, down: 697; *naïve vs. withdrawal*: 1039 DE transcripts, up: 510, down: 529) (Fig2A-B). Pathway enrichment analysis revealed that during intoxication, DE genes were associated with neurotransmission and addiction processes, including vesicle-mediated transport, synaptic signaling (glutamatergic synapse, dopaminergic synapse, oxytocin signaling pathways), NMDA receptor activation, and canonical addiction pathways (cocaine, nicotine, amphetamine) as well as circadian entrainment (Fig2F). During withdrawal, enriched pathways included inflammatory signaling (positive regulation of interleukin-8 production), stress responses (chaperone-mediated protein folding), cytoskeletal remodeling, and continued involvement of neurotransmitter signaling and morphine addiction pathways (Fig2G). Slightly fewer DE transcripts were found in the *naïve vs. abstinence* comparison (686 DE transcripts: 301 up, 385 down), suggesting a partial return toward baseline gene expression (Fig2C). However, pathways associated with cellular stress and organ repair (plasma membrane repair), synaptic signaling (GABAergic synapse), energy metabolism (glycolysis, NADH metabolism), and cytoskeletal remodeling remained altered (Fig2H).

**Figure 2.**
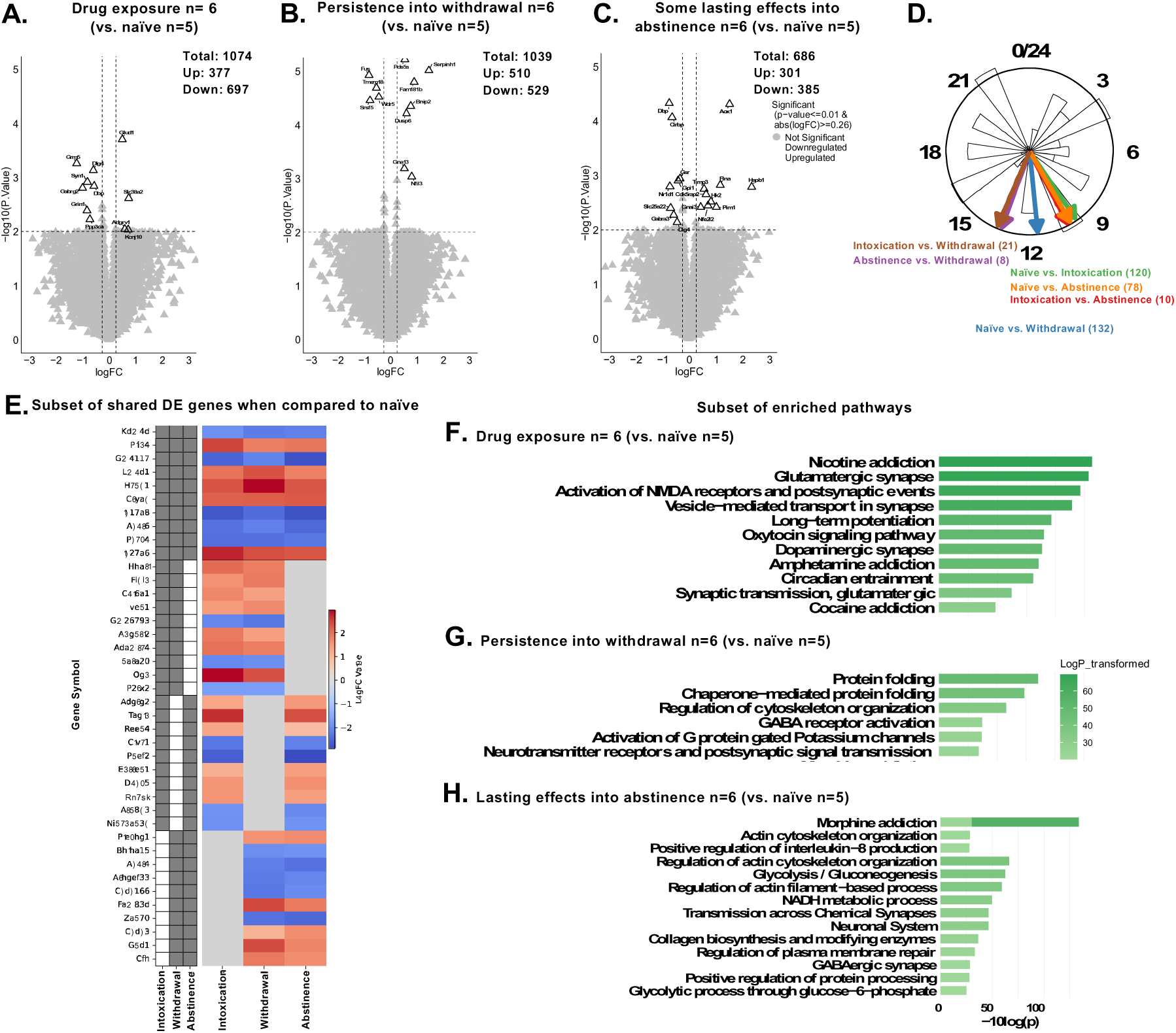
State-specific transcriptomic changes in the female SCN during opioid exposure, with lasting stress and repair signatures in abstinence. Volcano plots show differentially expressed (DE) genes in females comparing (A) naïve vs. intoxication, (B) naïve vs. withdrawal, and (C) naïve vs. abstinence. Genes with *p*<0.01 and |fold change|>0.26 are highlighted; upregulated genes are in red, downregulated in blue. (D) Polar plot depicts the average peak phase of rhythmic DE genes for each comparison. (E) Heatmap displays log_₂_ fold changes for genes DE in at least one drug-exposed group relative to naïve, with shared DE status indicated on the left (dark gray boxes). Selected enriched pathways are shown for (F) naïve vs. intoxication, (G) naïve vs. withdrawal, and (H) naïve vs. abstinence, with the x-axis representing –log10(*p*-value), where higher values indicate greater significance.

Comparisons between drug-exposed states revealed fewer differences (*intoxication vs. withdrawal*: 123 DE; *intoxication vs. abstinence*: 70 DE; *withdrawal vs. abstinence*: 38 DE), indicating that the most significant transcriptomic shifts occurred relative to the drug-naïve state.

A core subset of genes exhibited persistent regulation across multiple exposure states, indicating long-lasting molecular changes in the SCN. Genes involved in lipid metabolism (Plin4, Slc27a6) and cellular stress response (Hspb1, Cryab) remained consistently upregulated, suggesting disrupted energy homeostasis and sustained stress signaling (Fig2E). Additionally, several DE genes showed rhythmic expression profiles. Peak phases were detected for comparisons including *naïve vs. intoxication* (120 genes, CT10.15), *naïve vs. abstinence* (78 genes, CT9.71), *naïve vs. withdrawal* (132 genes, CT11.72), *intoxication vs. withdrawal* (21 genes, CT13.55), *abstinence vs. withdrawal* (8 genes, CT13.33), and *intoxication vs. abstinence* (10 genes, CT10) (Fig2D). These findings highlight temporally regulated, state-dependent rhythmic gene expression in females, with withdrawal-related comparisons showing a later phase. Overall, opioid exposure in female rats causes SCN transcriptomic changes, with acute effects during intoxication and withdrawal and lasting stress, metabolic, and circadian disruptions into abstinence. Metadata, gene expression counts, differential expression results, enriched pathways, and rhythmic DEGs across conditions are detailed in supplementary tables S1-S5.

### 3.2. Transcriptomic Changes in Males Reflect Acute Disruptions and Persistent Immune System Elevation

In male rats, a total of 1127 DE transcripts were identified, with a nearly equal distribution of upregulated (590) and downregulated (537) transcripts. Most changes occurred during intoxication (*naïve vs. intoxication*: 369 DE transcripts, up: 242, down: 127) and were enriched in pathways related to DNA damage response and cell cycle regulation (DNA damage checkpoint signaling), neurotransmission and synaptic plasticity (AMPA receptor trafficking, glutamate binding), immune regulation (response to type II interferon), and carbohydrate metabolism (hexose transmembrane transport) (Fig3A-F).

**Figure 3.**
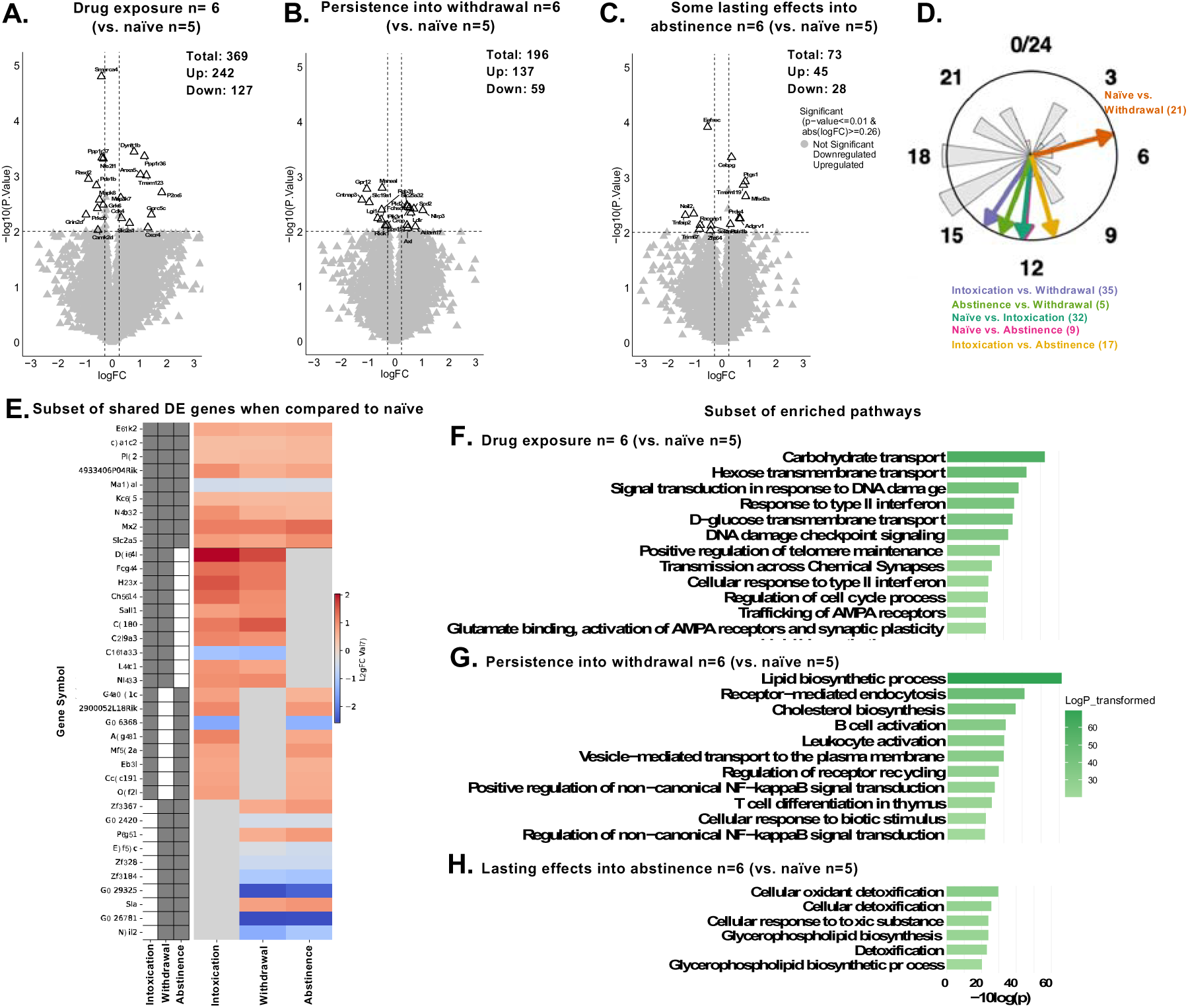
Transcriptomic changes in males across opioid exposure states, with persistent immune system activation. Volcano plots show differentially expressed (DE) genes in males comparing (A) naïve vs. intoxication, (B) naïve vs. withdrawal, and (C) naïve vs. abstinence. Genes with *p*<0.01 and |fold change|>0.26 are highlighted; upregulated genes are shown in red on the right, and downregulated genes are shown in blue on the left. (D) Polar plot depicting the average peak phase of rhythmic DE genes for each comparison. (E) Heatmap displaying log₂ fold changes for genes DE in at least one drug exposure group (intoxication, withdrawal, or abstinence) relative to naïve controls, with shared DE status shown on the left (dark gray boxes indicating significant DE). Gene expression levels are indicated by intensity, with red for upregulation and blue for downregulation. Selected enriched pathways are shown for (F) naïve vs. intoxication, (G) naïve vs. withdrawal, and (H) naïve vs. abstinence, with the x-axis representing –log10(*p*-value), where higher values correspond to more significant pathways.

Changes persisted into withdrawal (*naïve vs. withdrawal*: 196 DE transcripts, up: 137, down: 59; *intoxication vs. withdrawal*: 325 DE transcripts, up: 96, down: 229) with enriched pathways related to immune response and inflammation (B cell activation, T cell differentiation), lipid metabolism (cholesterol biosynthesis, vesicle-mediated transport), and cellular signaling (NF-kappaB signaling, cellular response to biotic stimuli) (Fig3B,G). These findings suggest that withdrawal is characterized by persistent disruptions in metabolism, cellular stress, and synaptic signaling, reflecting continued adaptation rather than a return to baseline. Slightly fewer changes were observed in abstinence (*intoxication vs. abstinence*: 107 DE transcripts, up: 41, down: 66; *withdrawal vs. abstinence*: 57 DE transcripts, up: 29, down: 28; *naïve vs. abstinence*: 73 DE transcripts, up: 45, down: 28), suggesting a partial return to baseline (Fig3C, H). However, pathways related to cellular detoxification and stress response (cellular oxidant detoxification, cellular detoxification, cellular response to toxic substance, detoxification) and lipid metabolism and membrane dynamics (glycerophospholipid biosynthesis, glycerophospholipid biosynthetic process) remained elevated. Most transcriptomic changes were most pronounced during intoxication and withdrawal, with trends toward baseline in abstinence, though immune-related pathways remained elevated, suggesting lasting effects.

A subset of genes was differentially expressed across all drug exposure states (intoxication, withdrawal, and abstinence), suggesting sustained transcriptional alterations in the SCN following chronic oxycodone exposure. These genes were associated with intracellular signaling (Pld2, Etnk2), protein regulation (Tceanc2, Nrbp2), and immune responses (Mx2). Notably, Slc2a5, a gene encoding the fructose transporter GLUT5, was dysregulated across all phases, pointing to altered metabolic sensing in the SCN. These persistent changes highlight stable shifts in SCN function that may contribute to long-term physiological consequences of opioid use in males (Fig3E).

A subset of DE genes also exhibited rhythmic expression patterns. Peak phases were observed for the following comparisons: *naïve vs. intoxication* (32 genes, CT12.33), *naïve vs. abstinence* (9 genes, CT12.44), *intoxication vs. abstinence* (17 genes, CT12.12), *intoxication vs. withdrawal* (21 genes, CT14.24), *abstinence vs. withdrawal* (9 genes, CT12.44), and *naïve vs. withdrawal* (32 genes, CT12.33) (Fig3D). Rhythmic expression was largely similar across groups, with the exception of the withdrawal vs. abstinence comparison, where rhythmic DE genes exhibited earlier peak phases. Overall, chronic opioid exposure in male rats leads to persistent transcriptomic changes in the SCN, with acute disruptions during intoxication and withdrawal, and lasting alterations in immune response, metabolism. Metadata, gene expression counts, differential expression results, enriched pathways, and rhythmic DEGs across conditions are detailed in supplementary tables S1-S5.

### 3.3. Distinct sex-specific transcripts emerged during abstinence, potentially regulated by rhythmic genes

A total of 337 DE transcripts were influenced by sex across all groups, with more downregulated (214) than upregulated (123) transcripts. Sex differences were observed in all drug states: 115 in naïve (up: 50, down: 65), 59 in intoxication (up: 17, down: 42), 70 in withdrawal (up: 32, down: 38), and 93 in abstinence (up: 24, down: 69) (Fig4A-D). Although the number of DE genes affected by sex was similar across groups, the specific sets of DE transcripts were distinct. Only three transcripts were shared across all states, with minimal overlap between pairs, such as naïve and withdrawal (5), naïve and abstinence (1), and abstinence and withdrawal (1) (Fig4L).

**Figure 4.**
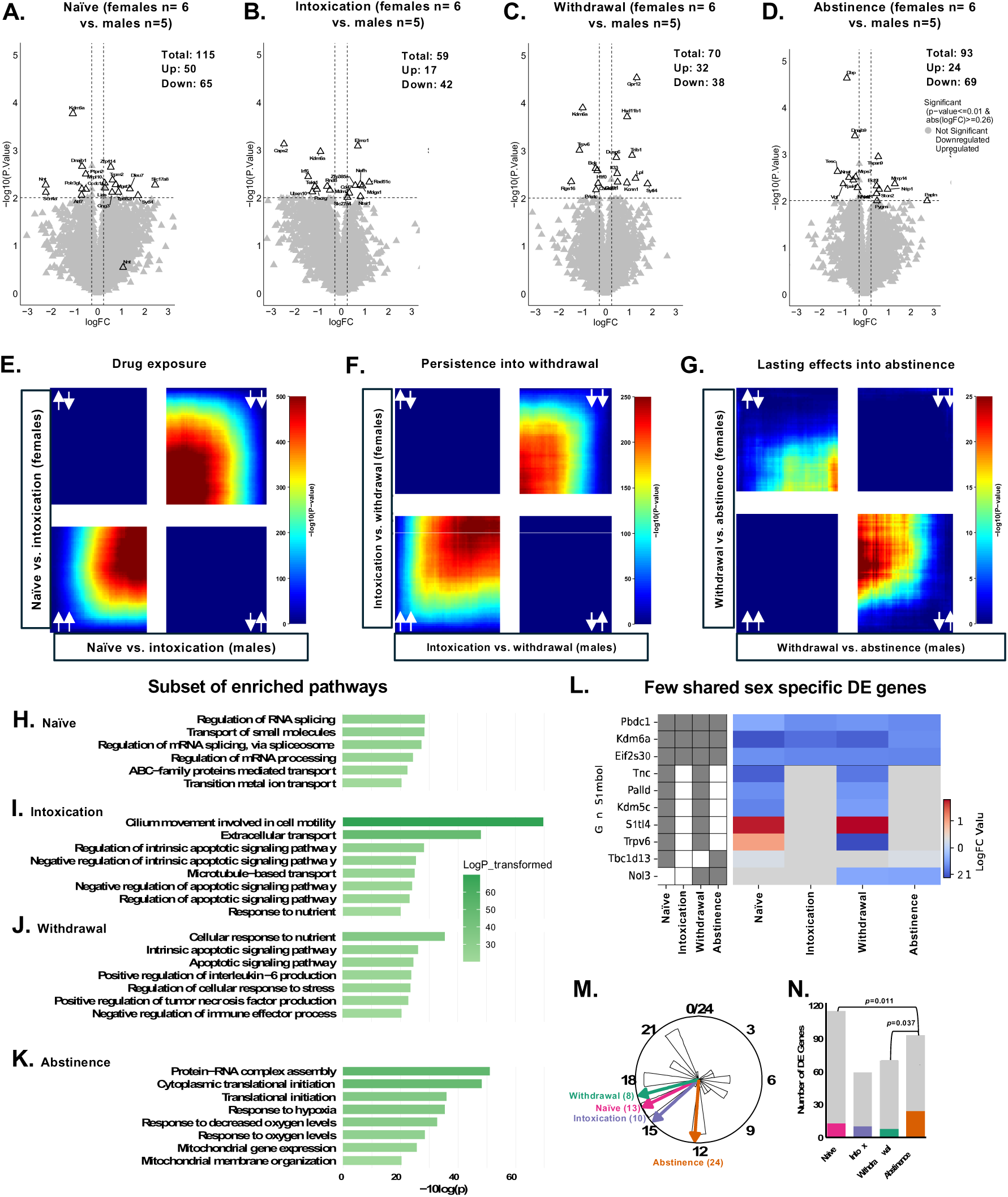
Similar numbers of sex-specific transcripts across drug states, with a divergent gene expression pattern emerging during abstinence. Volcano plots comparing sex differences in (A) naïve, (B) intoxication, (C) withdrawal, and (D) abstinence. Genes with p-value<0.01 and |FC|>0.26 are highlighted. Upregulated genes are shown on the right in red, while downregulated genes are shown on the left in blue. RRHO heatmap illustrating both downregulated genes (top-right), opposing regulation trends (top-left and bottom-right), and both upregulated genes (bottom-left), with warmer colors indicating higher -log10 p-values, highlighting sex specific patterns in (E) naïve vs. intoxication, (F) intoxication vs. withdrawal, and (G) withdrawal vs. abstinence. Subsets of enriched pathways are shown in (H) naïve, (I) intoxication, (J) withdrawal, and (K) abstinence, with the x-axis representing -10log(p-value) of each pathway, where higher values indicate more significant pathways. (L) Heatmap showing minimal overlap of sex-specific DE genes between groups. (M) Polar plot displaying the average peak phase time of rhythmic DE genes, and (N) a higher proportion of rhythmic DE genes to total DE genes in abstinence.

In the naïve state, sex differences were enriched in RNA processing and splicing (regulation of RNA splicing) and molecular transport (transition metal ion transport) (Fig4H). During intoxication, pathways related to cellular structure, motility, and transport (extracellular transport, microtubule-based transport), cell death and survival pathways (regulation of intrinsic apoptotic signaling pathway), and metabolic regulation (response to nutrient) were enriched (Fig4I). In withdrawal, enriched pathways focused on inflammation and immune regulation (positive regulation of interleukin-6 production, positive regulation of tumor necrosis factor production), apoptosis and cell death pathways (apoptotic signaling pathway), and cellular metabolism and stress response (cellular response to nutrient, regulation of cellular response to stress) (Fig4J). In abstinence, enriched pathways included cellular responses to stress, response to hypoxia, response to decreased oxygen levels, and mitochondrial function (mitochondrial gene expression, mitochondrial membrane organization) (Fig4K). The minimal overlap in DE transcripts and divergent enriched pathways suggest that sex modulates SCN function differently across drug states, leading to distinct molecular responses throughout addiction.

To further explore these differences, we applied RRHO—a threshold-free method to compare sex-specific gene expression patterns. Extensive overlap between sexes was found during intoxication (*naïve vs. intoxication*, Fig4E) and the transition into withdrawal (*intoxication vs. withdrawal*, Fig4F), while marked discordance emerged during the shift from withdrawal to abstinence (Fig4G), suggesting that sex-specific transcriptional patterns become more pronounced during abstinence.

A subset of sex-specific DE genes exhibited rhythmic expression profiles. The following peak phases were observed: naïve (13 genes, CT16), intoxication (10 genes, CT14.3), withdrawal (8 genes, CT15.8), and abstinence (24 genes, CT11.24) (Fig4M). The proportion of rhythmic DE genes was significantly higher in abstinence compared to both naïve (p=0.011) and withdrawal (p=0.037) phases (Fig4N), suggesting that these rhythmic genes could be key drivers of the sex differences observed in abstinence. Metadata, gene expression counts, differential expression results, enriched pathways, rhythmic DEGs, and concordant transcripts from RRHO analysis are provided in supplementary tables S1-S6.

### 3.4. Co-expression network analysis reveals distinct hub genes associated with intoxication

Weighted Gene Co-Expression Network Analysis (WGCNA) identified multiple gene modules significantly associated with intoxication, each characterized by distinct hub gene signatures and enriched pathways. The black module, enriched for genes involved in synaptic function, neuronal signaling, and intracellular transport, contained hub genes such as Vsnl1, Rbfox3, Syt7, Ache, and Rapgefl1, which are implicated in calcium signaling, synaptic vesicle cycling, and neurotransmitter regulation. Pathway enrichment analysis revealed significant involvement in *vesicle-mediated transport in synapse*, *synaptic vesicle cycle*, and neurotransmitter-specific signaling pathways including *dopamine*, *serotonin*, *glutamate*, and *opioid signaling.* Additionally, pathways related to circadian entrainment, long-term potentiation, and oxytocin signaling suggested broader alterations in neural plasticity. Interestingly, enrichment for cardiac-related pathways (cardiac conduction, calcium regulation in cardiac cells) pointed to potential peripheral or autonomic effects of opioid exposure (Fig5A-B). The red module, which contained genes related to cilia and microtubule-based structures such as Cfap52, Spag8, Tekt1, and Enkur, was enriched for pathways involving cilium- or flagellum-dependent motility, motor protein activity, sarcomere organization, and skeletal muscle contraction. This suggests a role for cytoskeletal remodeling and intracellular transport. Additionally, developmental pathways related to heart development and left/right asymmetry determination were enriched, highlighting the involvement of intoxication in activating genes linked to cellular architecture (Fig5C-D). The yellow module, consisting primarily of genes like Tubb2a, Rtn1, Rnf208, and Myo6, indicated elevated translational and cellular activity during intoxication. Enriched pathways included responses to oxidative stress, multicellular organismal stress, axon injury, and synaptic signaling (Fig5E-F). Additional details for the green, blue, and turquoise modules are shown in supplementary Fig1A-D.

**Figure 5.**
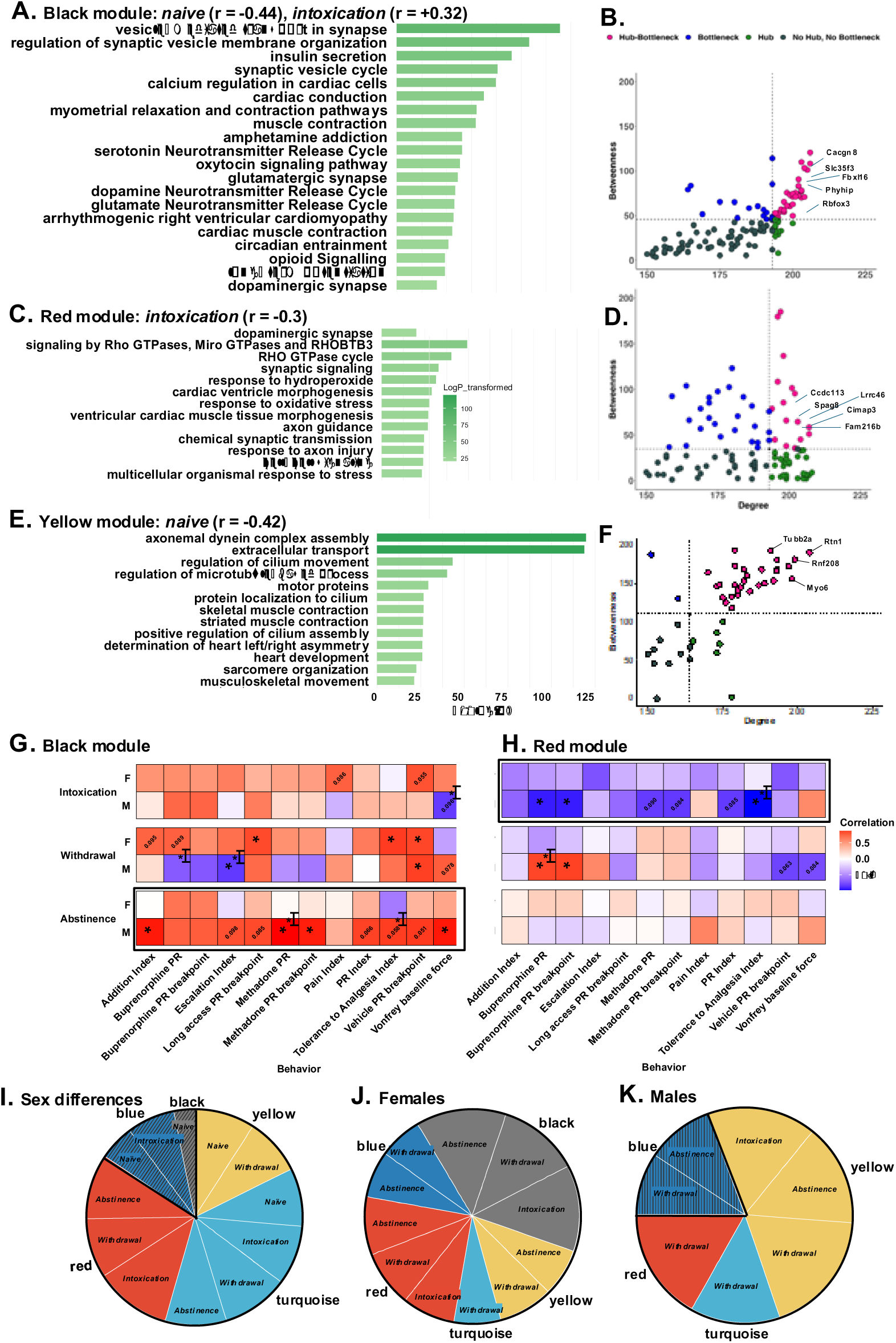
Co-expression analysis reveals intoxication-related hub genes and transcriptional signatures linked to addiction behavior. Enriched pathways for each identified module: (A) black, (C) red, and (E) green modules, with the x-axis representing the -log10 p-value to indicate statistical significance. Hub gene analysis for each module, identifying hub genes based on their high degree (number of connections to other genes) and betweenness centrality (extent to which a gene lies on the shortest paths between other genes) for (B) black, (D) red, and (F) yellow modules, respectively. Principal component analysis (PCA) was performed on gene expression data from the (G) black and (H) red modules, extracting PC1 as a summary measure, which was then correlated with behavioral indices to assess sex-specific relationships between gene expression and behavior across drug exposure states. Module differential connectivity analysis, conducted (I) sex-specifically, in (J) females only, and in (K) males only, reveals modules with gain or loss of transcriptional connectivity across drug states, with slice size proportional to the magnitude of connectivity change, solid colors indicating significant gain, and horizontal stripes indicating significant loss.

To link SCN gene co-expression with addiction behaviors, we summarized each WGCNA module using PC1 within each drug group and found distinct, group-dependent behavioral correlations. In the abstinence group, more severe addiction phenotypes were correlated with higher expression of black module genes, which are involved in synaptic signaling and neurotransmitter regulation (Fig5G). In the intoxication group, more severe addiction phenotypes were correlated with lower expression of red module genes, which are involved in cilia and microtubule function (Fig5H). These transcriptional signatures within specific SCN gene networks may underlie persistent alterations following opioid exposure and track with behavioral vulnerability. Additional behavior correlations are shown in supplementary Fig1E-F.

Module differential connectivity ratio (R) quantifies intramodular connectivity changes between drug states, with values >1 indicating increased connectivity and <1 suggesting decreased connectivity. Naïve females exhibited reduced connectivity in the black and blue modules (black: R=0.43, FDR<0.01; blue: R=0.68, FDR<0.01). In contrast, males showed increased connectivity in the red module across intoxication (R=1.46, FDR=0.012), abstinence (R=1.25, FDR<0.01), and withdrawal (R=1.14, FDR<0.01). The turquoise module also showed increased connectivity in males during intoxication (R=1.14, FDR=0.036), withdrawal (R=1.28, FDR<0.01), and abstinence (R=1.18, FDR<0.01), while the yellow module showed increased connectivity in naïve (R=1.14, FDR=0.012) and withdrawal (R=1.11, FDR<0.01) (Fig5I).

In females, the black and red modules exhibited increased connectivity across abstinence, intoxication, and withdrawal (black: Intoxication, R=2.46; Withdrawal, R=2.60; Abstinence, R=2.75; red: Intox, R=1.70; Withdrawal, R=1.75; Abstinence, R=1.59, all FDR<0.05). The blue and yellow modules showed increased connectivity in abstinent and withdrawal states (blue: Intoxication, R=1.15; Withdrawal, R=1.32; Abstinence, R=1.41; yellow: Withdrawal, R=1.65; Abstinence, R=1.43, all FDR<0.01). The turquoise module showed increased connectivity only in the withdrawal state (R=1.41, FDR<0.01) (Fig5J). In males, the blue module showed reduced connectivity in both abstinence (R=0.85, FDR=0.012) and withdrawal states (R=0.92, FDR<0.01). The yellow module exhibited increased connectivity across all drug states (Intoxication: R=1.41, FDR<0.01; Abstinence: R=1.52, FDR<0.01; Withdrawal: R=1.70, FDR<0.01). Both the magenta and red modules showed significant increases in connectivity during withdrawal (magenta: R=1.56, FDR=0.048; red: R=1.54, FDR=0.048). The turquoise module showed increased connectivity only in the withdrawal state (R=1.23, FDR<0.01) (Fig5K). Details on WGCNA module membership, hub and bottleneck genes, enriched pathways, PCA of gene expression, behavioral measures, and MDC are provided in supplementary tables S7–S12. Additional significant modules are shown in supplementary Fig1G-H.

## 4. Discussion

This study provides new insights into the sex-specific transcriptomic alterations within the SCN during opioid exposure and its relationship to addiction-like behaviors in genetically diverse heterogeneous stock (HS) rats. By examining the effects of oxycodone self-administration on SCN gene expression across four key phases (naïve, intoxication, withdrawal, and abstinence), we identified significant sex differences, which were particularly pronounced during abstinence, in both the acute and long-term effects of opioid use. These findings highlight the dynamic and persistent nature of SCN alterations in OUD and suggest that these changes contribute to the behavioral and physiological consequences of addiction.

Our data demonstrate that female and male rats exhibit distinct molecular responses in the SCN to opioid exposure. In females, intoxication and withdrawal were marked by changes in pathways related to neurotransmission, synaptic function, circadian rhythms, and inflammation. These alterations were accompanied by a lasting cellular stress-related transcriptional signature that persisted into abstinence. In contrast, males exhibited changes primarily related to immune regulation, DNA damage, and metabolism, with persistent pathways involved in immune system activation and cellular detoxification observed during abstinence. The sustained stress-related pathways observed in abstinence in females could reflect a heightened vulnerability to the physiological effects of opioid use, while the immune-related changes in males may underlie long-term neuroinflammatory responses following opioid exposure. These sex differences mirror prior reports of divergent opioid responses between males and females [39, 40]. While some studies suggest women have higher opioid craving [41, 42] and relapse risk during early abstinence [43], others report lower relapse rates after prolonged abstinence [44], and some find no sex differences [45, 46]. In rodents, females self-administered more oxycodone than males after acquisition of behavior [31] and showed higher reinstatement after short-access fentanyl and heroin self-administration [47, 48]. Findings on cue- and stress-induced reinstatement, as well as incubation of opioid craving, are mixed, with some studies reporting sex differences [49] and others not [50, 51]. These findings highlight the complex, sex-specific molecular adaptations to opioid exposure, suggesting that distinct mechanisms may underlie the differential vulnerability and long-term consequences of opioid use in males and females.

RRHO analysis reinforced this divergence, revealing more overlap between sexes in early drug states (intoxication and withdrawal) but marked discordance in abstinence. Notably, both sexes showed changes in sex-specific rhythmic DE genes, identified by cross-referencing DE genes with a known SCN-rhythmic gene database. Abstinence showed a significantly higher proportion of rhythmic, sex-dependent DE genes compared to intoxication and withdrawal, with the peak phases of these genes occurring earlier in abstinence than in the other groups. The earlier peak of sex-specific rhythmic DE genes during abstinence suggests a sex-dependent circadian contribution to opioid responses.

Co-expression network analysis revealed distinct modules of genes associated with different drug states, with the black and red module showing positive and negative associations with intoxication, respectively. To link SCN gene co-expression with addiction-related behaviors, we summarized each WGCNA module within each drug state and observed distinct, state-specific behavioral correlations. In the abstinence group, greater addiction severity was associated with elevated expression of genes involved in synaptic signaling and neurotransmitter regulation, suggesting that persistent neural signaling adaptations may contribute to long-term vulnerability. In contrast, during intoxication, more severe addiction phenotypes were linked to reduced expression of genes related to cilia and microtubule function, pointing to potential disruptions in cellular structure or signaling. Together, these findings suggest that specific SCN gene networks track with behavioral vulnerability in a drug group dependent manner and may underlie lasting molecular changes following opioid exposure.

The use of HS rats in this study is a strength, as it allows for a more accurate representation of the genetic diversity observed in human populations. This approach provides a more robust model for studying the complex, multifactorial nature of opioid addiction and its effects on the SCN, compared to inbred strains. Additionally, the inclusion of both male and female rats in our analysis provides a comprehensive understanding of the sex-specific molecular mechanisms underlying opioid addiction. Some limitations of our work include the use of bulk RNA sequencing, which does not allow for the identification of cell-type specific changes in gene expression, which may be important for understanding the cellular mechanisms by which opioids affect the SCN.

In summary, our findings demonstrate that opioid exposure induces sex- and state-specific transcriptomic changes in the SCN, which may contribute to behavioral and physiological outcomes in OUD. These findings underscore the SCN as a critical brain region involved in the pathophysiology of OUD. Future studies should investigate how these molecular alterations in the SCN interact with other brain regions involved in addiction, and whether targeting circadian pathways could provide a novel approach for treating OUD.

## 5. Acknowledgements None

## 6. Author contributions

OG, AAP, and RWL obtained funding for the project. RWL conceptualized and managed the project and provided supervision for data analyses and interpretations. TCD, SS, and RWL analyzed datasets. LCSW, AAP, OG, and RWL provided expertise on rat experiments. LM and OG provided rat tissues. BRW and MCG processed rat tissues and prepared tissues for sequencing. TCD and RWL wrote the manuscript, created the figures, supplemental information. All authors participated in reviewing and editing the manuscript for publication.

---

**Supplementary Fig 1.**
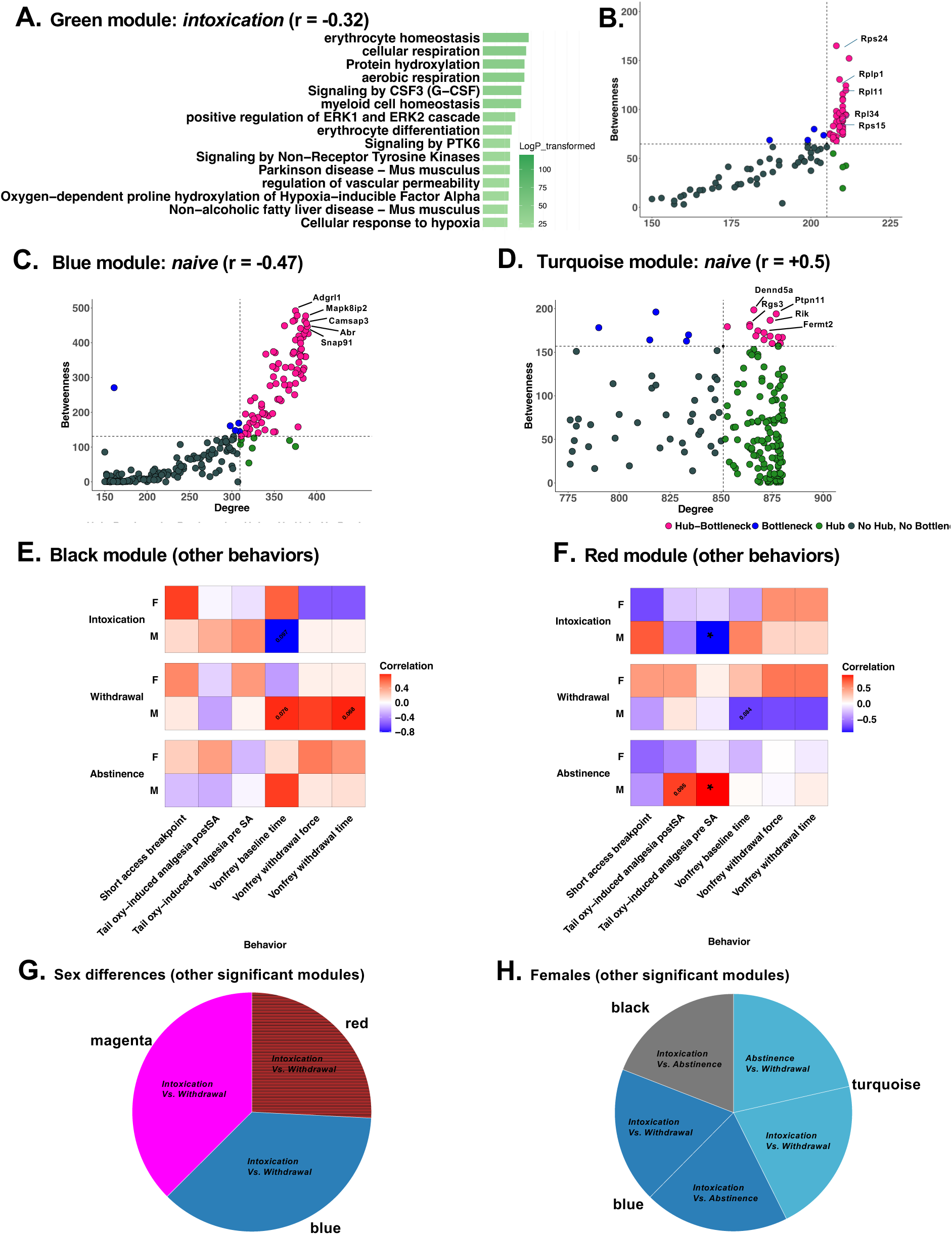

## References

1. Keyes, K.M., et al., What is the prevalence of and trend in opioid use disorder in the United States from 2010 to 2019? Using multiplier approaches to estimate prevalence for an unknown population size. Drug and Alcohol Dependence Reports, 2022. 3: p. 100052.

2. Kenan, K., K. Mack, and L. Paulozzi, Trends in prescriptions for oxycodone and other commonly used opioids in the United States, 2000-2010. Open Med, 2012. 6(2): p. e41–7.

3. McQuay, H., Opioids in pain management. The Lancet, 1999. 353(9171): p. 2229–2232.

4. Sanger, N., et al., Treatment Outcomes in Patients With Opioid Use Disorder Who Were First Introduced to Opioids by Prescription: A Systematic Review and Meta-Analysis. Frontiers in Psychiatry, 2020. 11.

5. Information, C.I.f.H. Pan-Canadian trends in theprescribing of opioids 2012 to 2016. 2017.

6. Prevention, C.f.D.C.a. US opioid prescribing rate maps. Atlanta: Centers for Disease Control and Prevention 2018; Available from: https://www.cdc.gov/drugoverdose/maps/rxrate-maps.html.

7. Abuse, N.I.o.D. As Opioid Use Disorders Increased, Prescriptions for Treatment Did Not Keep Pace. 2020; Available from: https://www.drugabuse.gov/news-events/nida-notes/2018/07/opioid-use-disorders-increased-prescriptions-treatment-did-not-keep-pace.

8. Lankenau, S.E., et al., Initiation into prescription opioid misuse amongst young injection drug users. International Journal of Drug Policy, 2012. 23(1): p. 37–44.

9. Administration, S.A.a.M.H.S. Results from the 2019 National Survey on Drug Use and Health: Detailed Tables. 2020; Prevalence Estimates, Standard Errors, P Values, and Sample Sizes.].

10. Prevention, C.f.D.C.a. Understanding Drug Overdoses and Deaths. 2023; Available from: https://archive.cdc.gov/www_cdc_gov/drugoverdose/epidemic/index.html.

11. Gossop, M., et al., Lapse, relapse and survival among opiate addicts after treatment: A prospective follow-up study. The British Journal of Psychiatry, 1989. 154(3): p. 348–353.

12. Mahfoud, Y., et al., Sleep disorders in substance abusers: how common are they? Psychiatry (Edgmont), 2009. 6(9): p. 38.

13. Cao, M. and S. Javaheri, Effects of chronic opioid use on sleep and wake. Sleep medicine clinics, 2018. 13(2): p. 271–281.

14. Shaw, I.R., et al., Acute intravenous administration of morphine perturbs sleep architecture in healthy pain-free young adults: a preliminary study. Sleep, 2005. 28(6): p. 677–682.

15. Howe, R.C., F.W. Hegge, and J.L. Phillips, Acute heroin abstinence in man: II. Alterations in rapid eye movement (REM) sleep. Drug and alcohol dependence, 1980. 6(3): p. 149–161.

16. Shi, J., et al., Long-term methadone maintenance reduces protracted symptoms of heroin abstinence and cue-induced craving in Chinese heroin abusers. Pharmacology Biochemistry and Behavior, 2007. 87(1): p. 141–145.

17. Brower, K.J., et al., PER3 polymorphism and insomnia severity in alcohol dependence. Sleep, 2012. 35(4): p. 571–577.

18. O’Connor, P.G. and D.A. Fiellin, Pharmacologic treatment of heroin-dependent patients. Treatment of Substance Use Disorders, 2013: p. 87–114.

19. Von Korff, M., et al., Long-Term Opioid Therapy Reconsidered. Annals of Internal Medicine, 2011. 155(5): p. 325–328.

20. Roehrs, T.A. and T. Roth, Sleep Disturbance in Substance Use Disorders. Psychiatr Clin North Am, 2015. 38(4): p. 793–803.

21. Kay, D.C., Human sleep during chronic morphine intoxication. Psychopharmacologia, 1975. 44(2): p. 117–124.

22. Dimsdale, J.E., et al., The effect of opioids on sleep architecture. Journal of clinical sleep medicine, 2007. 3(01): p. 33–36.

23. Zhang, R., et al., Rest-Activity Rhythms, Their Modulators, and Brain-Clinical Correlates in Opioid Use Disorder. JAMA Network Open, 2025. 8(2): p. e2457976.

24. Xue, X., et al., Molecular rhythm alterations in prefrontal cortex and nucleus accumbens associated with opioid use disorder. Translational Psychiatry, 2022. 12(1).

25. Puig, S., et al., Sex specific role of the circadian transcription factor NPAS2 in opioid tolerance, withdrawal and analgesia. Genes, Brain and Behavior, 2022. 21(7).

26. Baird, T.J. and D.V. Gauvin, Characterization of Cocaine Self-Administration and Pharmacokinetics as a Function of Time of Day in the Rat. Pharmacology Biochemistry and Behavior, 2000. 65(2): p. 289–299.

27. Bass, C.E., H.T. Jansen, and D.C.S. Roberts, FREE-RUNNING RHYTHMS OF COCAINE SELF-ADMINISTRATION IN RATS HELD UNDER CONSTANT LIGHTING CONDITIONS. Chronobiology International, 2010. 27(3): p. 535–548.

28. Du, K., et al., Melatonin attenuates fentanyl - induced behavioral sensitization and circadian rhythm disorders in mice. Physiology & Behavior, 2024. 279: p. 114523.

29. Shelton, M.A., et al., Sex-specific Concordance of Striatal Transcriptional Signatures of Opioid Addiction in Human and Rodent Brains. Biological Psychiatry Global Open Science, 2025: p. 100476.

30. Kallupi, M., et al., Nociceptin attenuates the escalation of oxycodone self-administration by normalizing CeA–GABA transmission in highly addicted rats. Proceedings of the National Academy of Sciences, 2020. 117(4): p. 2140–2148.

31. Kimbrough, A., et al., Oxycodone self-administration and withdrawal behaviors in male and female Wistar rats. Psychopharmacology, 2020. 237: p. 1545–1555.

32. Mavrikaki, M., et al., Oxycodone self-administration in male and female rats. Psychopharmacology, 2017. 234: p. 977–987.

33. Carrette, L.L.G., et al., The Cocaine and Oxycodone Biobanks, Two Repositories from Genetically Diverse and Behaviorally Characterized Rats for the Study of Addiction. eneuro, 2021. 8(3): p. ENEURO.0033-21.

34. Barko, K. and et al., Brain region- and sex-specific transcriptional profiles of microglia. Frontiers in Psychiatry, 2022. 13.

35. Ritchie, M.E., et al., limma powers differential expression analyses for RNA-sequencing and microarray studies. Nucleic acids research, 2015. 43(7): p. e47–e47.

36. Cahill, K.M., et al., Improved identification of concordant and discordant gene expression signatures using an updated rank-rank hypergeometric overlap approach. Scientific Reports, 2018. 8(1): p. 9588.

37. Zhang, R., et al., A circadian gene expression atlas in mammals: implications for biology and medicine. Proceedings of the National Academy of Sciences, 2014. 111(45): p. 16219–16224.

38. Panda, S., et al., Coordinated transcription of key pathways in the mouse by the circadian clock. Cell, 2002. 109(3): p. 307–20.

39. Nicolas, C., et al., Sex differences in opioid and psychostimulant craving and relapse: a critical review. Pharmacological reviews, 2022. 74(1): p. 119–140.

40. Knouse, M.C. and L.A. Briand, Behavioral sex differences in cocaine and opioid use disorders: The role of gonadal hormones. Neuroscience & Biobehavioral Reviews, 2021. 128: p. 358–366.

41. Back, S.E., et al., Comparative profiles of men and women with opioid dependence: results from a national multisite effectiveness trial. Am J Drug Alcohol Abuse, 2011. 37(5): p. 313–23.

42. Yu, J., et al., Gender and stimulus difference in cue-induced responses in abstinent heroin users. Pharmacology Biochemistry and Behavior, 2007. 86(3): p. 485–492.

43. Maehira, Y., et al., Factors associated with relapse into drug use among male and female attendees of a three-month drug detoxification-rehabilitation programme in Dhaka, Bangladesh: a prospective cohort study. Harm Reduct J, 2013. 10: p. 14.

44. Gordon, M.S., et al., A randomized clinical trial of buprenorphine for prisoners: Findings at 12-months post-release. Drug Alcohol Depend, 2017. 172: p. 34–42.

45. Herbeck, D.M., et al., Gender differences in treatment and clinical characteristics among patients receiving extended release naltrexone. J Addict Dis, 2016. 35(4): p. 305–314.

46. Kennedy, A.P., et al., Sex differences in cocaine/heroin users: drug-use triggers and craving in daily life. Drug Alcohol Depend, 2013. 132(1-2): p. 29–37.

47. Malone, S.G., et al., Escalation and reinstatement of fentanyl self-administration in male and female rats. Psychopharmacology (Berl), 2021. 238(8): p. 2261–2273.

48. Smethells, J.R., et al., Effects of voluntary exercise and sex on multiply-triggered heroin reinstatement in male and female rats. Psychopharmacology (Berl), 2020. 237(2): p. 453–463.

49. Vazquez, M., et al., Acute ovarian hormone treatment in freely cycling female rats regulates distinct aspects of heroin seeking. Learn Mem, 2020. 27(1): p. 6–11.

50. Phillips, A.G., et al., Oral prescription opioid-seeking behavior in male and female mice. Addict Biol, 2020. 25(6): p. e12828.

51. Reiner, D.J., et al., Relapse to opioid seeking in rat models: behavior, pharmacology and circuits. Neuropsychopharmacology, 2019. 44(3): p. 465–477.

