## Supplemental Tables for "Sex-specific transcriptional signatures of oxycodone persist during withdrawal and abstinence in the suprachiasmatic nucleus of heterogeneous stock rats"

Table S1. Metadata

Table S2. Gene Expression Counts

Table S3. DEG with Ensembl ID, Log2 Fold Change, and p-values

Table S4. Enriched Pathways from Metascape Analysis of DEG

Table S5. Rhythmic DEG Identified Across Conditions

Table S6. Lists of concordant transcripts identified with RRHO

Table S7. Weighted Gene Co-Expression Network Analysis (WGCNA) Module Membership

Table S8. WGCNA-Identified Hub and Bottleneck Genes by Module

Table S9. Enriched Pathways from Metascape of Genes in WGCNA Modules

Table S10. Principal Component Analysis of Gene Expression

Table S11. Behavioral Measures and Index Scores

Table S12. Module Differential Connectivity Results from WGCNA
